## Supplementary Information for "Mapping the glycosyltransferase fold landscape using deep learning"

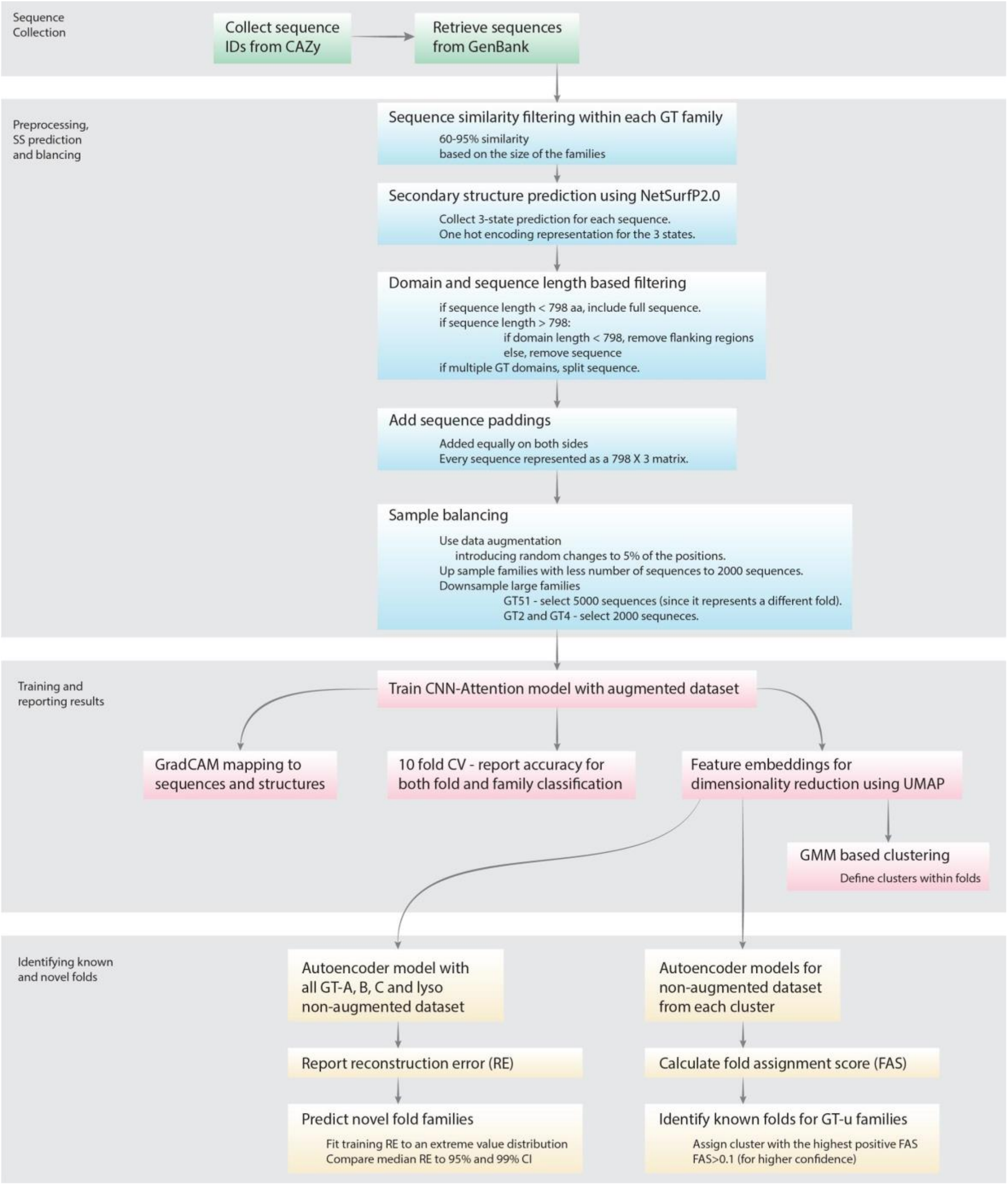


**Supplementary Figure 1: Flowchart showing the preprocessing, training and interpretation steps of the CNN-Attention and autoencoder model.**

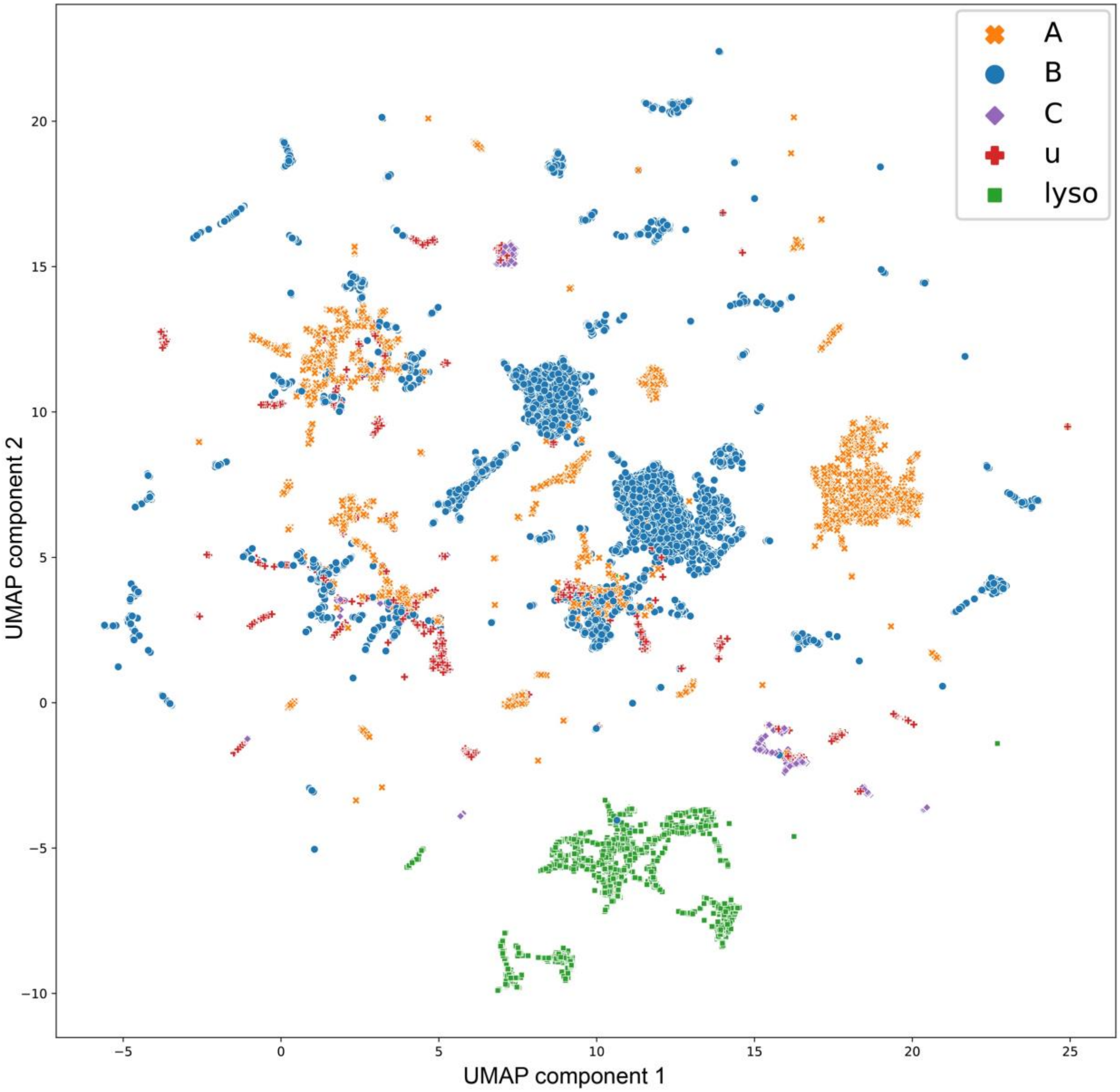


**Supplementary Figure 2: Projection of Transformer based model embeddings.** The projection is generated from the embeddings of all trained GT sequences obtained using the transformer based esm-1b method. Following esm-1b instructions, the mean values are taken across all positions to generate a vector of 1280 dimensions. UMAP was applied to this high dimensional data to generate these two dimensional projections for visualization of GT fold clusters. While projection representations for sequences of same fold generally group together, the obtained groups are not as well segregated as in the CNN-attention model.

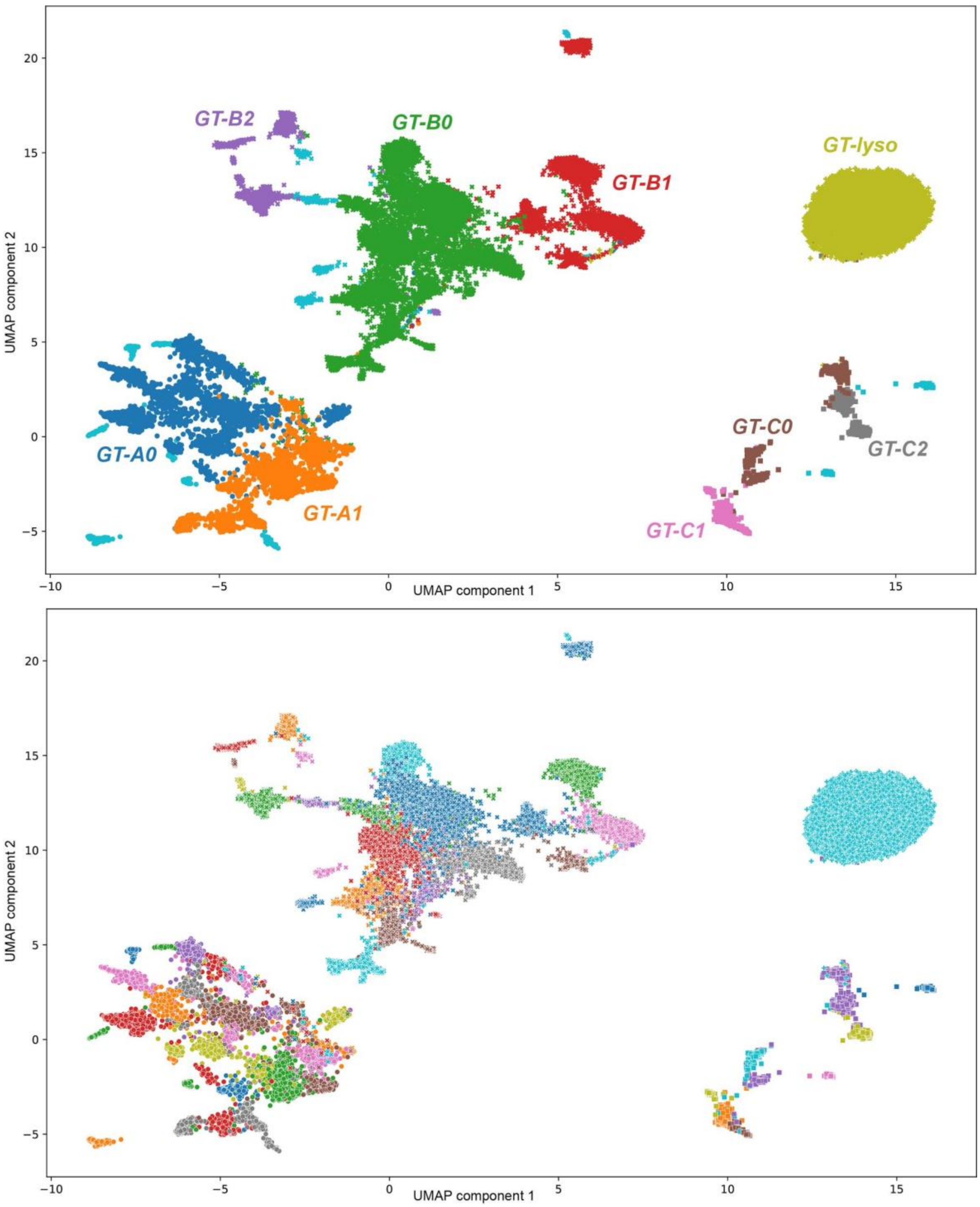


**Supplementary Figure 3: UMAP projections for the feature vectors of the major GT fold types.** Sequences are colored based on their cluster assignments (Top) and based on CAZy families (Bottom). Both plots show sequences from the same cluster and family grouping together, respectively.

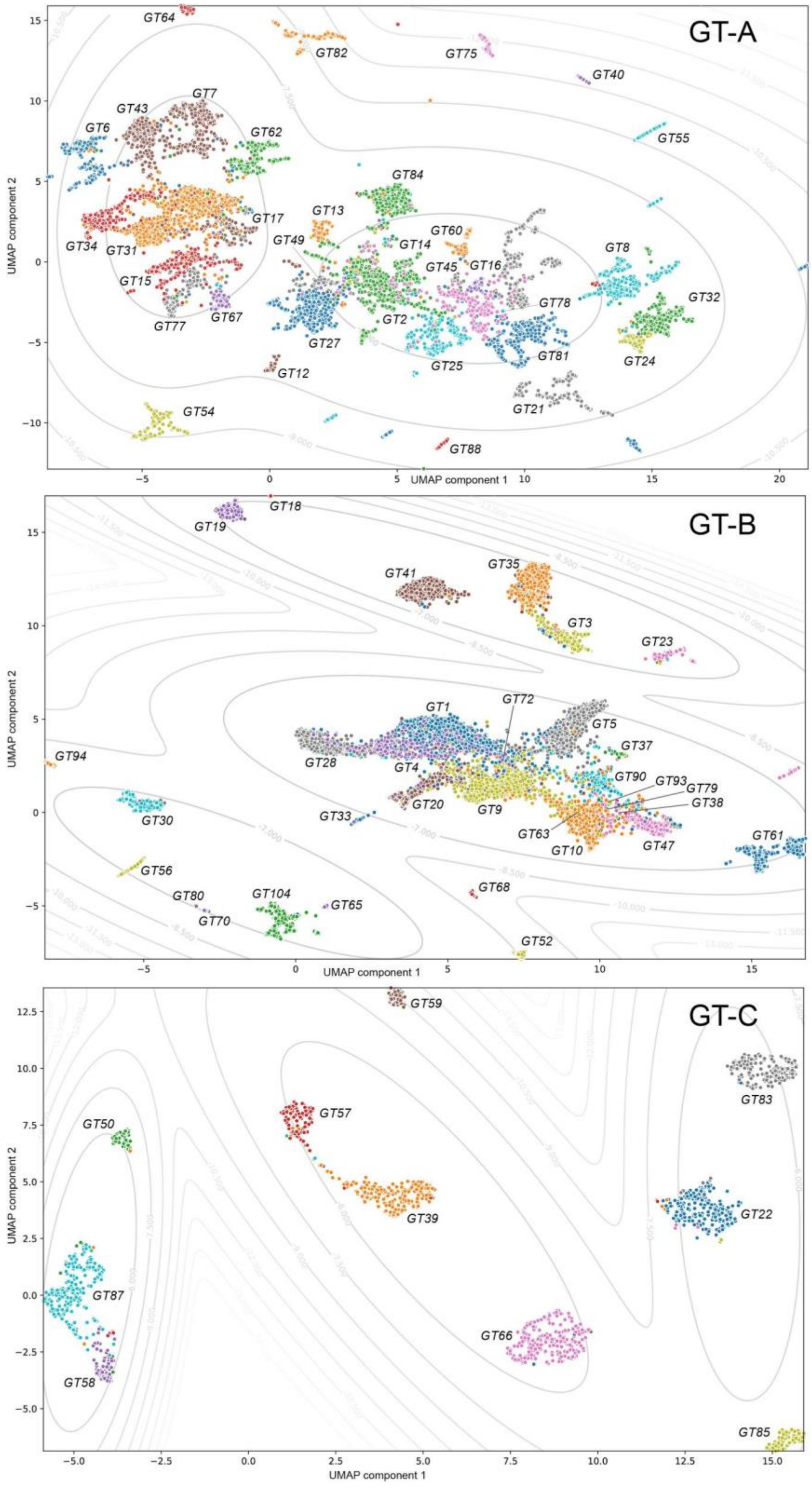


**Supplementary Figure 4: UMAP projections for the feature vectors from separate fold types.** Sequences are colored based on their family and labelled.

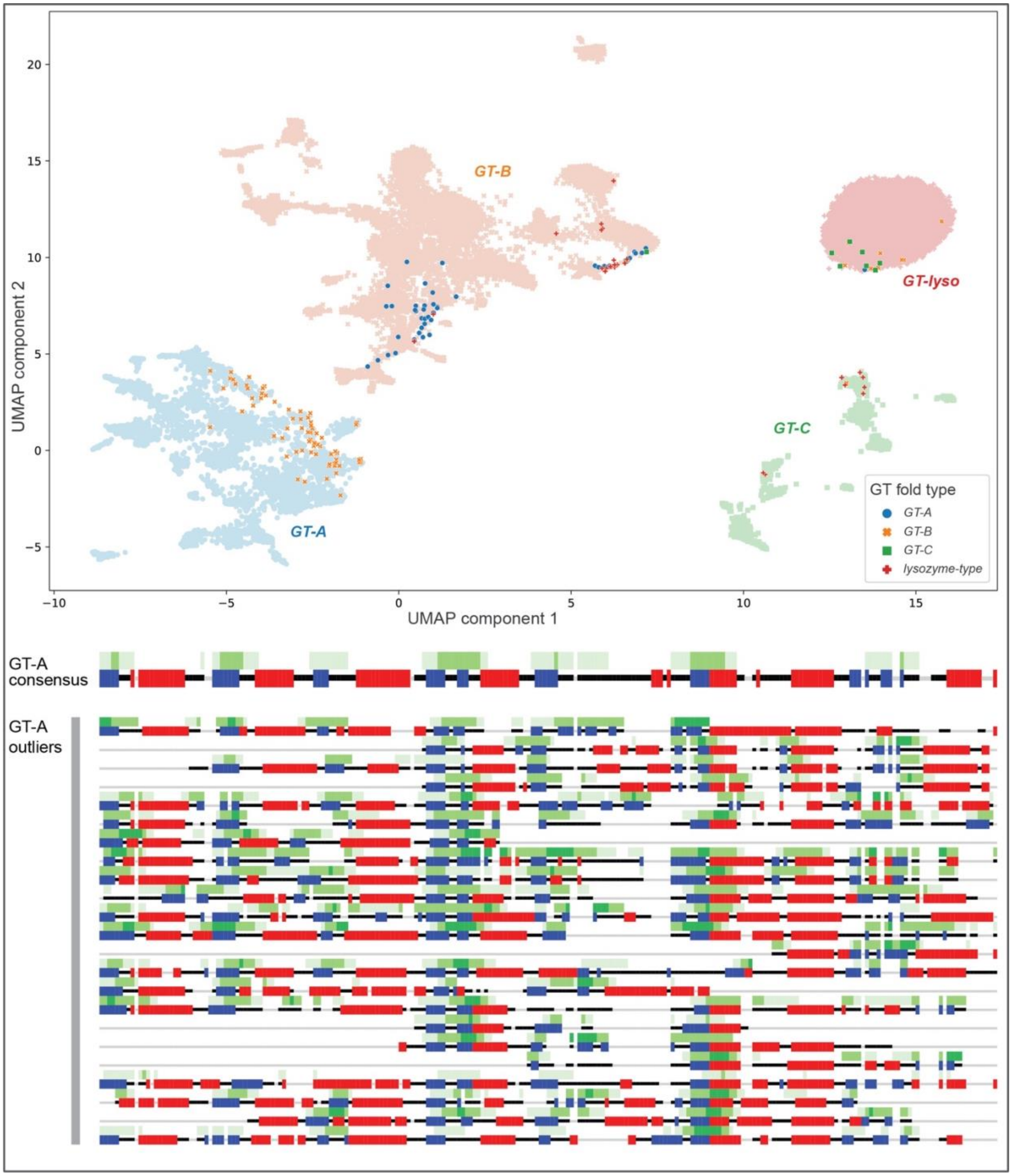


**Supplementary Figure 5: Analysis of the outlier sequences.** Top: Outlier sequences are highlighted in the UMAP projections. Most of these sequences include fragmentary sequences. Bottom: Layer 2 CAM maps for the aligned GT-A domain of the outlier sequences compared to the GT-A consensus. Other folds not shown since no consensus alignment for comparison.

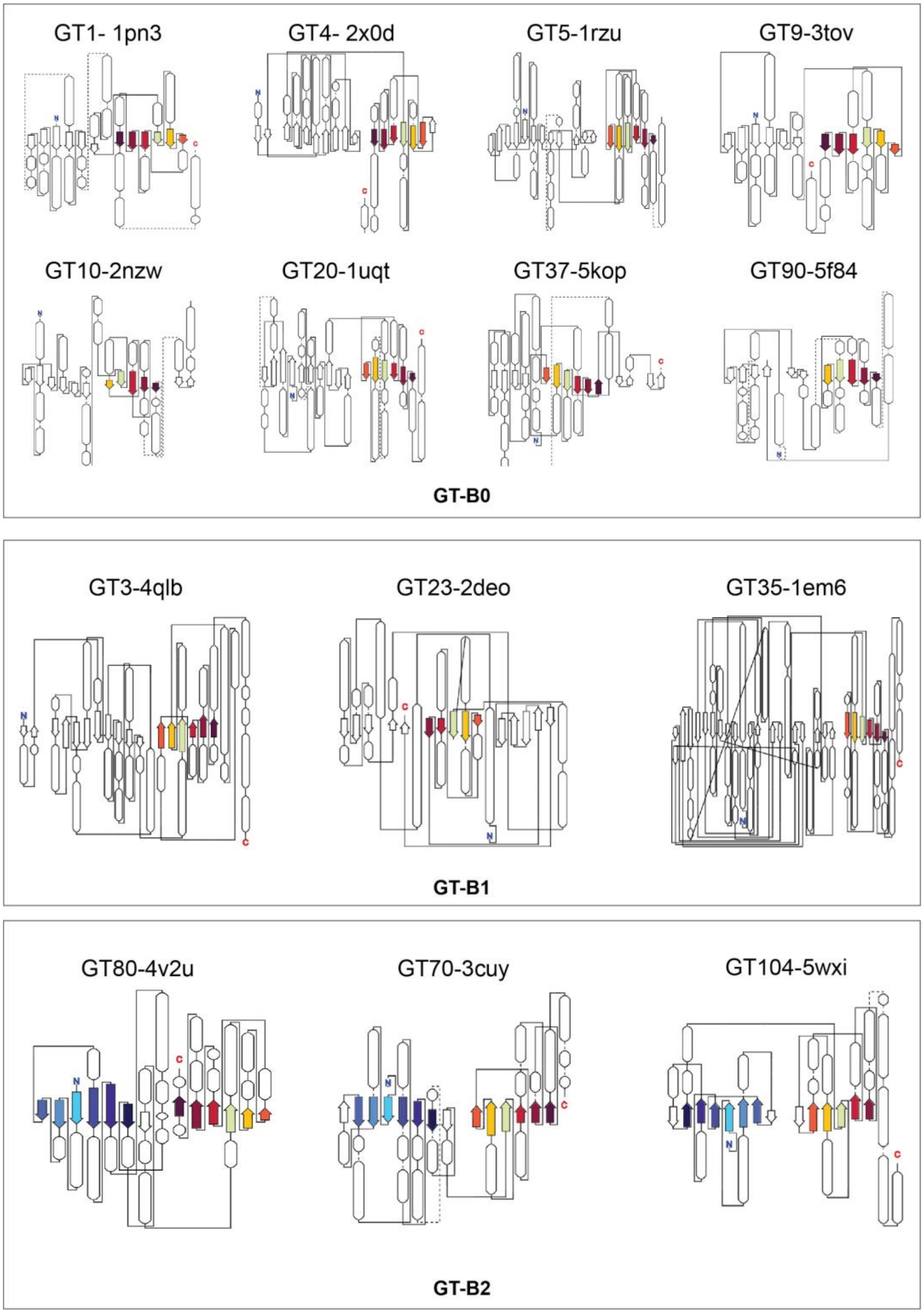


**Supplementary Figure 6: Topologies for multiple representative structures for GT-B families from different clusters show the core conserved features identified by the CNN-Attention module.**

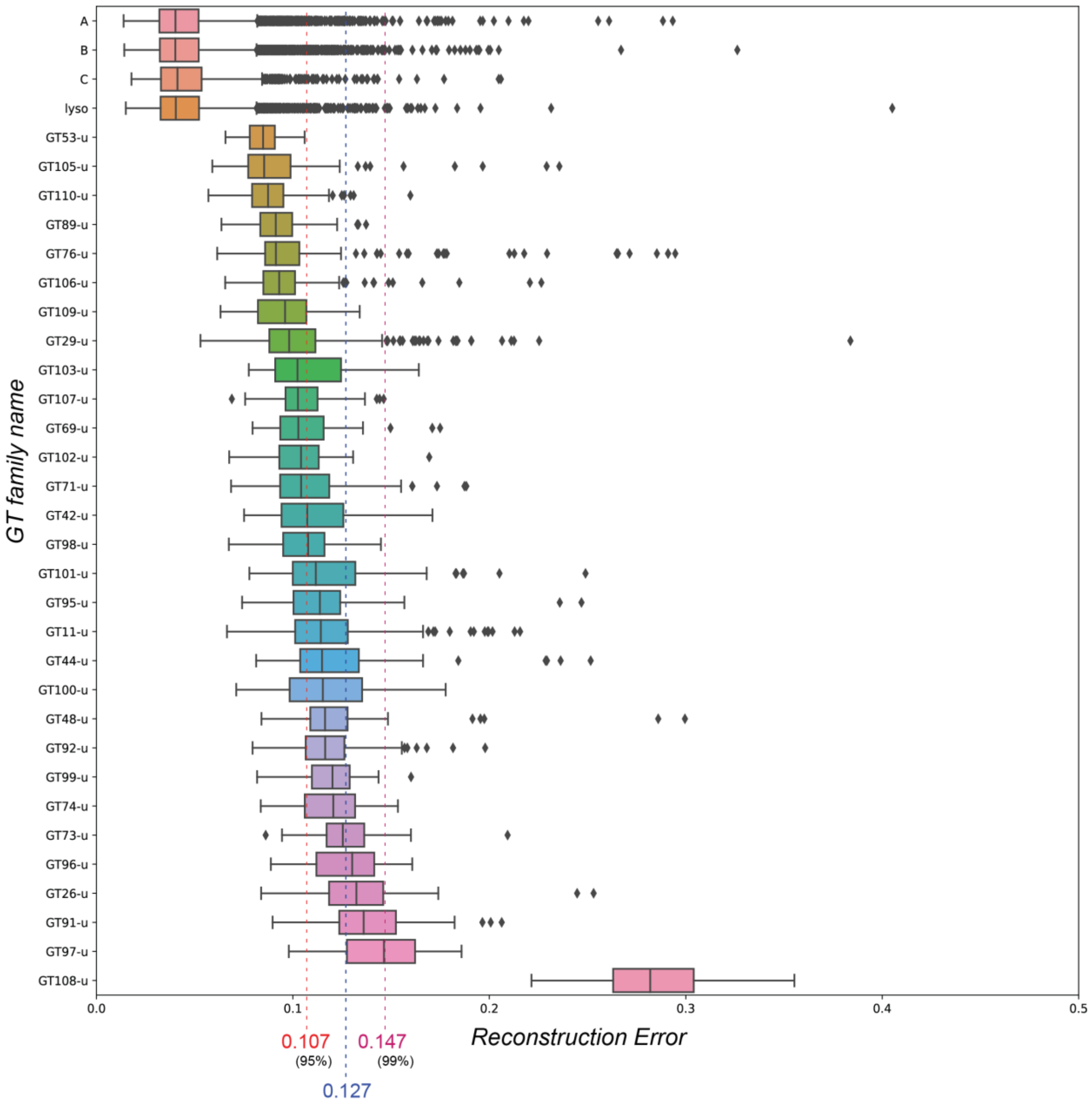


**Supplementary Figure 7: Boxplot showing the RE for each of the 4 known GT folds and all the GT-u families.** Red line at 0.104 indicates the 95%CI upper bound based on the extreme value distribution of the training (known GT folds) sequences. Magenta line at 0.147 indicates the 99% CI upper bound. Blue line at 0.127 indicates the mid-point. GT-u families with median RE above this value are predicted to have novel GT folds with increasing confidence.

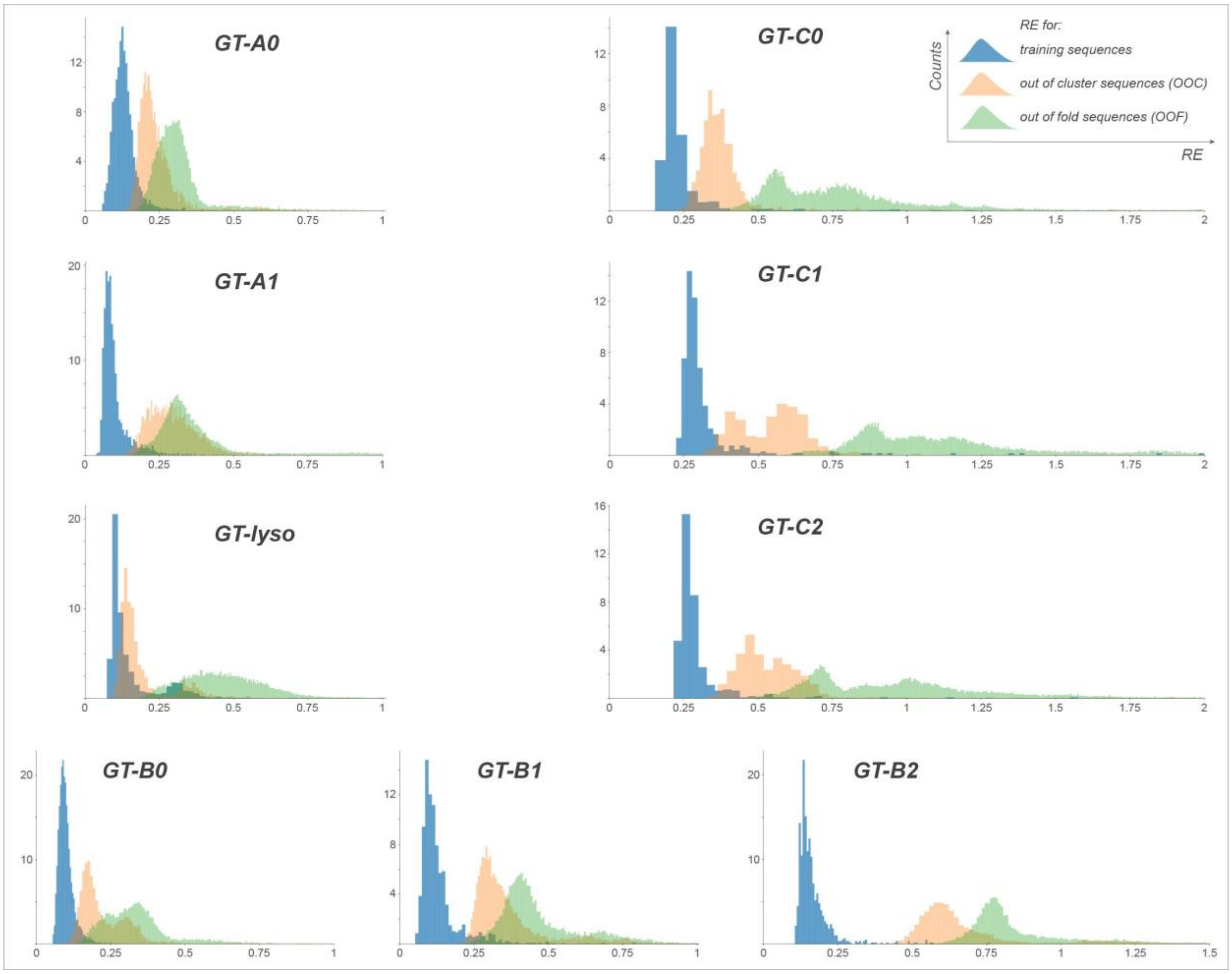


**Supplementary Figure 8: RE distribution for the training, out of cluster and out of fold sequences for each of the 9 clusters generated using the cluster specific autoencoder models.**

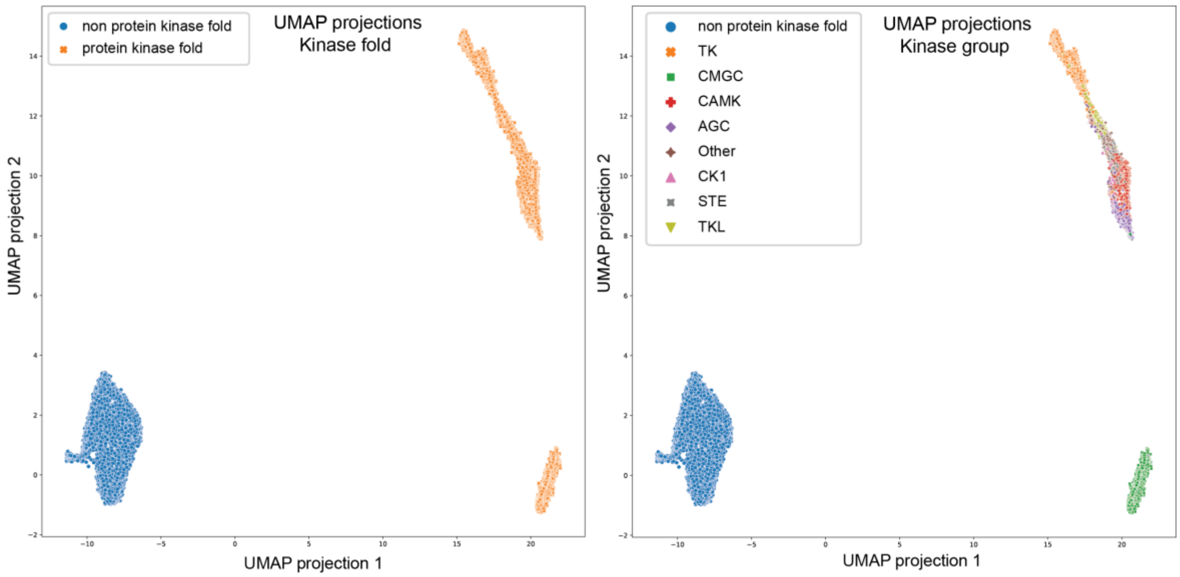


**Supplementary Figure 9: UMAP projections obtained from the CNN-attention model trained on 11675 kinase sequences.** The 2D projections show a clear separation of protein kinase fold sequences from non-protein kinase fold sequences (left). Separation also places kinases from the same group together (right). The accuracy for protein kinase fold separation from non-protein kinase fold is 99%. The accuracy for the separation of kinase groups is 83%.

**Supplementary Table 1: A complete comparison of different modules in the CNN-attention model.** The model was trained with the same hyperparameter settings: learning rate at 5e-5 and weight decay rate at 1e-5 using Adam optimizer. Datasets were separated as augmented (Aug) or non-augmented (Non-Aug) to make comparison of the effect of the data augmentation method. The effects of multitask learning and attention modules were also tested.

| Model | Dataset | Target | Precision | Recall | Accuracy | F1-score |
|---|---|---|---|---|---|---|
| CNN+ Multitask | Non-Aug | Fold | 0.85 | 0.87 | 0.87 | 0.86 |
| | | Family | 0.23 | 0.29 | 0.29 | 0.23 |
| CNN+ Multitask+ Attention | Non-Aug | Fold | 0.90 | 0.90 | 0.90 | 0.89 |
| | | Family | 0.29 | 0.35 | 0.35 | 0.29 |
| CNN+ Multitask | Aug | Fold | 0.92 | 0.92 | 0.92 | 0.92 |
| | | Family | 0.59 | 0.57 | 0.57 | 0.55 |
| CNN+ Attention | Aug | Fold | 0.96 | 0.96 | 0.96 | 0.96 |
| | | Family | 0.72 | 0.69 | 0.69 | 0.69 |
| CNN+ Multitask+ Attention | Aug | Fold | 0.96 | 0.96 | 0.96 | 0.96 |
| | | Family | 0.78 | 0.77 | 0.77 | 0.77 |

**Supplementary Table 2: Comparison of the prediction results from the GT CNN-attention model and the HHsearch results.** The best 3 hits from the HHsearch results with an e-value lower than 1e-2 is considered for fold assignment using HHsearch. Whether HHsearch results match the CNN-attention network and/or the results from previous literature is also indicated.

| GT-u fam | GT CNN-attention prediction | Literature support | HHsearch | Does HHsearch results match GT-CNN and literature support? | HHsearch results | | | | | |
|---|---|---|---|---|---|---|---|---|---|---|
| | | | | | 1st best hit | E-val | 2nd best hit | E-val | 3rd best hit | E-val |
| GT100-u | variant | - | GT-B | No | GT52-B | 6.21E-11 | GT38-B | 1.41E-08 | GT80-B | 1.00E-07 |
| GT101-u | variant | variant | GT-A | No | GT2-A | 7.06E-16 | GT45-A | 2.08E-13 | GT8-A | 8.14E-11 |
| GT102-u | GT-B | - | none | No | GT5-B | 2.52E-01 | GT5-B | 4.49E-01 | GT63-B | 5.22E-01 |
| GT103-u | GT-B | - | Both A and B | No | GT25-A | 9.45E-11 | GT4-B | 1.61E-06 | GT4-B | 5.93E-02 |
| GT105-u | GT-C | GT-C | GT-B | No | GT41-B | 3.12E-30 | GT41-B | 5.91E-25 | GT41-B | 1.40E-19 |
| GT106-u | GT-B | - | GT-B | Yes | GT65-B | 3.89E-06 | GT68-B | 1.26E-04 | GT23-B | 1.38E-04 |
| GT107-B | GT-B | - | GT-B | Yes | GT19-B | 3.57E-03 | GT4-B | 3.76E-02 | GT41-B | 1.11E-01 |
| GT108-u | novel | novel | none | Yes | GT4-B | 3.62E+00 | GT56-B | 4.44E+00 | GT35-B | 5.86E+00 |
| GT109-u | GT-A | GT-A | GT-A | Yes | GT54-A | 7.28E-21 | GT25-A | 3.09E-06 | GT13-A | 2.65E-03 |
| GT110-u | GT-A | - | GT-B | No | GT47-B | 1.19E-06 | GT10-B | 3.89E-03 | GT5-B | 1.70E-01 |
| GT11-u | variant | - | GT-B | No | GT23-B | 1.60E-10 | GT68-B | 6.31E-04 | GT65-B | 8.56E-04 |
| GT26-u | novel | novel | none | Yes | GT5-B | 1.03E-02 | GT41-B | 6.27E-01 | GT30-B | 1.14E+00 |
| GT29-u | GT-A | GT-A variant | none | No | GT56-B | 8.01E+00 | GT52-B | 9.20E+00 | GT20-B | 1.03E+01 |
| GT42-u | variant | GT-A variant | none | Yes | GT70-B | 1.71E+00 | GT57-C | 1.94E+00 | GT52-B | 2.42E+00 |
| GT44-u | variant | - | GT-A | No | GT32-A | 2.58E-11 | GT88-A | 3.55E-04 | GT63-B | 2.76E-01 |
| GT48-u | variant | - | none | Yes | GT10-B | 3.10E+00 | GT27-A | 8.52E+00 | GT33-B | 8.93E+00 |
| GT53-u | GT-C | GT-C | GT-C | Yes | GT83-C | 3.33E-03 | GT87-C | 4.88E-03 | GT87-C | 5.30E-02 |
| GT69-u | GT-A | - | GT-A | Yes | GT62-A | 3.06E-05 | GT15-A | 8.09E-01 | GT60-A | 1.02E+00 |
| GT71-u | GT-A | - | GT-A | Yes | GT8-A | 1.71E-03 | GT24-A | 1.10E-01 | GT32-A | 8.28E-01 |
| GT73-u | variant | - | none | Yes | GT57-C | 1.54E+00 | GT33-B | 6.92E+00 | GT16-A | 7.37E+00 |

| | | | | | | | | | | |
|---|---|---|---|---|---|---|---|---|---|---|
| GT74-u | variant | - | GT-A | No | GT2-A | 7.00E-21 | GT45-A | 1.25E-17 | GT27-A | 9.35E-17 |
| GT76-u | GT-C | - | GT-C | Yes | GT39-C | 2.55E-05 | GT50-C | 6.59E-04 | GT83-C | 1.77E-03 |
| GT89-u | GT-C | - | GT-C | Yes | GT83-C | 1.86E-03 | GT66-C | 1.96E-02 | GT39-C | 3.02E-01 |
| GT91-u | novel | - | none | Yes | GT75-A | 3.37E+00 | GT16-A | 5.89E+00 | GT20-B | 1.00E+01 |
| GT92-u | GT-A | - | none | No | GT6-A | 1.60E+00 | GT17-A | 2.01E+00 | GT55-A | 2.51E+00 |
| GT95-u | GT-A | - | none | No | GT6-A | 2.23E+00 | GT16-A | 2.81E+00 | GT65-B | 3.33E+00 |
| GT96-u | novel | - | none | Yes | GT6-A | 5.24E-02 | GT34-A | 1.68E-01 | GT8-A | 3.72E-01 |
| GT98-u | GT-C | - | GT-C | Yes | GT66-C | 1.22E-07 | GT6-A | 3.03E+00 | GT32-A | 5.63E+00 |
| GT99-u | variant | variant | GT-B | No | GT19-B | 3.00E-02 | GT25-A | 1.01E-01 | GT30-B | 2.58E-01 |

**Supplementary Table 3: A comparison of CNN-attention model with other state of the art models.** The comparison is performed on same training dataset as CNN-attention model. Three compared methods are Transformer embeddings with GBDT classifier, single layer LSTM model, ProtCNN model. The results are analysis based on four aspects, fold level accuracy, family level accuracy, interpretability and the ability to classify GT-u families. The interpretability and ability to handle unknown fold sequences were also compared across models. The CNN-attention model has interpretable outputs for every step compared to the Transformer+GBDT model which provides some clustering results based on the projections but does not provide any information about core conserved features, while the LSTM and ProtCNN methods do not provide any such interpretable outputs. Because we incorporate autoencoder models, GT CNN-attention is able to handle unknown fold sequences, whereas other methods cannot.

| **Model** | **Fold Accuracy** | **Family Accuracy** | **Interpretability** | **Classify GT-u** |
|---|---|---|---|---|
| **CNN-attention(ours)** | **0.96** | **0.77** | **Yes** | **Yes** |
| Transformer+GBDT | 0.96 | 0.67 | Limited | No |
| LSTM | 0.47 | 0.34 | No | No |
| ProtCNN | 0.84 | 0.76 | No | No |

**Supplementary Table 4: List of GT families and their corresponding fold and cluster.** A Gaussian Mixture Model (GMM) was used to cluster families based on their 2D UMAP projections generated separately for each fold type. Families with a GMM score above -7.5 for GT-A, -7 for GT-B and -6.5 for GT-C were placed in a cluster.

| Fold | Cluster | Families | GMM-Score |
|---|---|---|---|
| **GT-A** | **GT-A0** | GT16 | -5.4641 |
| | | GT2 | -5.524542 |
| | | GT60 | -5.642016 |
| | | GT14 | -5.688812 |
| | | GT45 | -5.694006 |
| | | GT25 | -5.785815 |
| | | GT78 | -5.786725 |
| | | GT49 | -5.945889 |
| | | GT21 | -6.060102 |
| | | GT27 | -6.150723 |
| | | GT84 | -6.30725 |
| | | GT13 | -6.346881 |
| | | GT24 | -6.38189 |
| | | GT81 | -6.477958 |
| | | GT8 | -6.500215 |
| | | GT32 | -6.599175 |
| | | GT12 | -7.134441 |
| | **GT-A1** | GT31 | -5.309125 |
| | | GT15 | -5.494145 |
| | | GT17 | -5.566161 |
| | | GT7 | -5.657536 |
| | | GT77 | -5.716002 |
| | | GT43 | -5.777749 |
| | | GT34 | -5.940145 |
| | | GT67 | -5.945407 |
| | | GT62 | -6.155893 |
| | | GT6 | -6.906416 |
| | **GT-A-Ungrouped** | GT88 | -7.635158 |
| | | GT64 | -8.277095 |
| | | GT54 | -8.51051 |
| | | GT82 | -9.230959 |
| | | GT55 | -9.308303 |
| | | GT40 | -10.256429 |
| | | GT75 | -11.761891 |
| **GT-B** | **GT-B0** | GT9 | -4.539272 |
| | | GT90 | -4.614016 |
| | | GT72 | -4.620983 |
| | | GT93 | -4.639714 |
| | | GT1 | -4.824178 |
| | | GT4 | -4.83733 |
| | | GT63 | -4.837775 |
| | | GT79 | -4.847818 |
| | | GT38 | -4.904147 |
| | | GT10 | -5.109297 |
| | | GT20 | -5.238405 |

| | | | |
|---|---|---|---|
| | | GT37 | -5.267101 |
| | | GT28 | -5.505519 |
| | | GT47 | -5.912927 |
| | | GT5 | -5.96545 |
| | | GT61 | -6.830745 |
| | **GT-B1** | GT41 | -5.310586 |
| | | GT3 | -5.656573 |
| | | GT35 | -5.760154 |
| | | GT23 | -6.101421 |
| | | GT19 | -6.834921 |
| | **GT-B2** | GT65 | -6.139725 |
| | | GT104 | -6.484054 |
| | | GT30 | -6.673493 |
| | | GT56 | -6.829993 |
| | | GT80 | -6.840005 |
| | | GT70 | -6.889586 |
| | **GT-B-Ungrouped** | GT33 | -7.144054 |
| | | GT18 | -7.231676 |
| | | GT94 | -7.454821 |
| | | GT52 | -8.463694 |
| | | GT68 | -8.781483 |
| **GT-C** | **GT-C0** | GT39 | -5.025598 |
| | | GT66 | -5.590182 |
| | | GT57 | -5.851775 |
| | **GT-C1** | GT87 | -4.413031 |
| | | GT58 | -5.296014 |
| | | GT50 | -6.05887 |
| | **GT-C2** | GT22 | -5.371296 |
| | | GT83 | -5.807555 |
| | **GT-C-Ungrouped** | GT85 | -6.935044 |
| | | GT59 | -8.834369 |
| **GT-lyso** | **GT-lyso** | GT51 | - |

**Supplementary Table 5: Fold prediction results for the GT-u families.** RE against all known GTs are shown. RE < 0.107 (Upper 95% CI) suggesting families most likely to adopt a known GT fold are highlighted in yellow and RE > 0.127 (Closer to or more than upper 99% CI) for families most likely to adopt a novel fold are highlighted in red. GT-u families are most likely to adopt the GT fold with the highest positive FAS. Families predicted to adopt a variant or a novel fold have negative FAS scores for all the clusters. The highest FAS scores for families predicted to adopt known folds are colored in green. The predicted fold and confidence are indicated in the last two columns. Confidence is evaluated based on the RE and FAS scores.

| Family | Reconstruction Error RE (against All known GTs) | Fold Assignment Scores (FAS) | | | | | | | | | | Prediction Confidence | Predicted fold |
|---|---|---|---|---|---|---|---|---|---|---|---|---|---|
| | | GT-A0 | GT-A1 | GT-B0 | GT-B1 | GT-B2 | GT-C0 | GT-C1 | GT-C2 | GT-lyso | Max FAS score | | |
| GT53-u | 0.0848 | -7.7062 | -17.1023 | -20.4061 | -3.4586 | -11.7955 | 1.6505 | 7.7203 | 2.6355 | -29.5924 | 7.7203 | High | GT-C |
| GT105-u | 0.0850 | -17.5652 | -24.5877 | -27.2568 | -6.0943 | -17.2715 | -3.7203 | 2.2301 | -10.6613 | -43.8454 | 2.2301 | High | GT-C |
| GT110-u | 0.0874 | 0.5617 | 2.1336 | 1.6885 | -2.1761 | 0.8386 | -28.8364 | -41.8105 | -26.1506 | -10.5905 | 2.1336 | High | GT-A |
| GT89-u | 0.0913 | -1.6840 | -4.6917 | -5.6962 | 0.8086 | -3.2460 | 1.7864 | 2.8052 | 6.1896 | -13.9826 | 6.1896 | High | GT-C |
| GT76-u | 0.0914 | -19.0593 | -26.4973 | -29.2158 | -5.7049 | -20.1115 | -4.4119 | 2.8857 | -10.5468 | -36.8289 | 2.8857 | High | GT-C |
| GT106-u | 0.0930 | -0.8764 | 0.2566 | 1.0686 | -0.3207 | -0.8118 | -19.1842 | -27.1105 | -15.6718 | -10.9195 | 1.0686 | High | GT-B |
| GT109-u | 0.0960 | 0.1085 | 0.5548 | 0.2431 | 0.2316 | -1.3323 | -20.6968 | -30.3417 | -19.6562 | -14.0250 | 0.5548 | Medium | GT-A |
| GT29-u | 0.0981 | -1.7044 | 0.4057 | -0.0964 | -4.1254 | -3.9635 | -34.3220 | -44.1672 | -32.0038 | -15.6071 | 0.4057 | Medium | GT-A |
| GT103-u | 0.1023 | -0.9478 | -0.3071 | -0.1846 | 0.2727 | 0.5684 | -17.7835 | -25.6814 | -10.9217 | -11.8482 | 0.5684 | Medium | GT-B |
| GT107-u | 0.1026 | -1.7008 | -0.3479 | 1.0348 | 0.1481 | 1.5433 | -20.0356 | -24.7787 | -12.7447 | -9.4429 | 1.5433 | Medium | GT-B |
| GT69-u | 0.1028 | -0.2249 | 1.3810 | -1.5294 | -3.4850 | -3.3949 | -30.2525 | -42.1769 | -27.9236 | -11.6703 | 1.3810 | Medium | GT-A |
| GT102-u | 0.1041 | -2.8304 | -2.1481 | 0.8966 | -1.3502 | -0.2873 | -32.6477 | -42.2431 | -28.8707 | -23.9149 | 0.8966 | Medium | GT-B |
| GT71-u | 0.1042 | -0.6198 | 0.5227 | -1.1886 | -0.4666 | -1.2484 | -16.2671 | -24.2920 | -11.2713 | -7.6293 | 0.5227 | Medium | GT-A |
| GT98-u | 0.1077 | -3.5928 | -7.0913 | -9.1323 | -0.5953 | -4.5628 | 1.3219 | 2.3093 | 6.5837 | -21.9022 | 6.5837 | Low | GT-C |
| GT92-u | 0.1164 | -1.0359 | 0.6344 | -4.7917 | -7.1016 | -5.0746 | -37.4749 | -49.0855 | -32.7090 | -13.6905 | 0.6344 | Low | GT-A |
| GT95-u | 0.1138 | -1.6167 | 0.2050 | -2.2480 | -4.9121 | -6.2884 | -35.9600 | -49.1779 | -31.2102 | -15.7293 | 0.2050 | Low | GT-A |
| GT42-u | 0.1073 | -2.3182 | -1.9045 | -0.9173 | -2.9590 | -5.5915 | -34.2709 | -47.3138 | -33.2359 | -21.2766 | -0.9173 | Low | Variant |
| GT101-u | 0.1117 | -3.4019 | -2.5153 | -0.7002 | -2.5644 | -4.1142 | -35.4129 | -47.4813 | -30.8002 | -26.3136 | -0.7002 | Low | Variant |
| GT11-u | 0.1143 | -3.9747 | -3.8743 | -1.6070 | -5.9780 | -7.9233 | -49.7788 | -64.5324 | -46.9072 | -35.2588 | -1.6070 | Medium | Variant |
| GT44-u | 0.1148 | -4.4052 | -7.1966 | -3.8032 | -1.8888 | -9.5213 | -27.6446 | -36.4394 | -27.4815 | -31.9549 | -1.8888 | Medium | Variant |
| GT100-u | 0.1152 | -4.2930 | -3.2511 | -1.1603 | -5.2216 | -5.5395 | -46.6134 | -60.9087 | -41.1213 | -31.2184 | -1.1603 | Medium | Variant |
| GT48-u | 0.1164 | -2.1581 | -4.3775 | -4.5470 | -0.0394 | -4.3777 | -8.9187 | -14.8223 | -4.6367 | -16.7056 | -0.0394 | Medium | Variant |
| GT99-u | 0.1201 | -4.0847 | -3.5176 | -0.4866 | -2.8529 | -4.2945 | -37.0718 | -47.9242 | -36.3760 | -28.2846 | -0.4866 | High | Variant |
| GT74-u | 0.1206 | -1.4762 | -0.6952 | -3.8812 | -5.7720 | -6.6236 | -39.2722 | -58.2878 | -35.5920 | -21.4848 | -0.6952 | High | Variant |
| GT73-u | 0.1254 | -3.3259 | -1.7022 | -2.3812 | -6.0193 | -8.2526 | -46.8867 | -61.8775 | -42.3077 | -27.5473 | -1.7022 | High | Variant |
| GT96-u | 0.1302 | -2.0410 | -0.3352 | -3.1068 | -2.1614 | -2.6451 | -22.3939 | -29.5370 | -17.4108 | -11.4877 | -0.3352 | Low | Novel |
| GT26-u | 0.1323 | -4.7941 | -7.0916 | -0.9723 | -5.9388 | -8.4612 | -43.8379 | -61.4936 | -45.9031 | -40.3355 | -0.9723 | Low | Novel |
| GT91-u | 0.1360 | -2.2260 | -0.9271 | -7.7585 | -6.2706 | -5.0966 | -31.1590 | -36.7761 | -23.2723 | -7.6507 | -0.9271 | Medium | Novel |
| GT97-u | 0.1464 | -7.6683 | -6.8926 | -1.3011 | -7.7251 | -8.8101 | -62.2334 | -77.3602 | -63.1995 | -44.5282 | -1.3011 | Medium | Novel |
| GT108-u | 0.2819 | -12.5744 | -11.9952 | -27.6863 | -34.7669 | -32.2872 | -94.3019 | -131.0731 | -93.7474 | -66.5097 | -11.9952 | High | Novel |
